# The role of microbial diversity in the formation of soil organic matter quality and persistence

**DOI:** 10.1101/2021.04.23.441131

**Authors:** Luiz A. Domeignoz-Horta, Melissa Shinfuku, Pilar Junier, Simon Poirier, Eric Verrecchia, David Sebag, Kristen M DeAngelis

## Abstract

The largest terrestrial carbon sink on earth is soil carbon stocks. As the climate changes, the rate at which the Earth’s climate warms depends in part on the persistence of soil organic carbon. Microbial turnover forms the backbone of soil organic matter (SOM) formation and it has been recently proposed that SOM molecular complexity is a key driver of stability. Despite this, the links between microbial diversity, chemical complexity and biogeochemical nature of soil organic matter remain missing. Here we used a model soil system to test the hypothesis that more diverse microbial communities generate more stable soil organic matter. We inoculated microbial communities of varying diversities into an model soil matrix amended with simple carbon, and measured the thermal stability of the resultant soil organic matter. Using a novel data analysis approach with Rock-Eval^®^ ramped thermal analysis, we found that microbial community diversity drives the chemical fingerprint of soil organic matter. Bacteria-only and low diversity communities lead to less chemically-diverse and more thermally-labile soil carbon pools than highly diverse communities. Our results provide direct evidence for a link between microbial diversity, molecular complexity and SOM stability. This evidence demonstrates the benefits of managing soils for maximum biological diversity as a means of building persistent SOM stocks.

**Classification:** Biological Sciences: Ecology

## Introduction

One of the grand challenging questions in microbiology is: when and where “who’s there” matters for ecosystem functioning^1^? It has been postulated that microbial community structure matters for phylogenetically “narrow” processes such as denitrification^2^, but not so much for phylogenetically-“broad” processes, such as carbon (C) cycling, which are completed by the majority of community members. However, recent work brings into question the assumption that all steps of carbon cycling are independent of microbial community composition^3,4^. Soil microbes are diverse in their macromolecular structures and metabolites^5^ and therefore microbial-derived SOM may reflect distinctions across communities. Recently, it was also hypothesized that molecular-diverse soil organic matter persists longer in soil^7^. Here we provide empirical data to support the hypothesis that distinct communities inoculated into a model soil shape the composition of soil organic matter, and that SOM formed under more diverse communities is more persistent to decomposition.

## Results & Discussion

Soil-derived microbial communities were subject to diversity removal by treatments with dilution (DO > D1 > D2), filtering (bacteria predominantly “B_*only*_”), and heat (spore forming “SF”), and incubated under different moisture and temperature in order to generate distinct microbial communities in an artificial soil matrix^4^. In a sibling study aiming to disentangle the drivers of carbon use efficiency, we observed that abiotic factors impact the microbial community assembly and characteristics, including Fungal:Bacterial ratio and biotic-driven soil aggregation^4^. However, it was the microbial community characteristics, e.g. bacterial community structure, bacterial richness, fungi presence, and extracellular enzymatic activity that impacted microbial carbon use efficiency^4^. Here, we analyzed the newly formed soil organic matter after four months of growth on cellobiose, using a method commonly used to quantify thermal stability and gradual stabilization of SOM^7^. The hydrocarbon compounds released at each temperature for each sample during the pyrolytic phase of Rock-Eval^®^ was used to calculate the Bray-Curtis-based chemical dissimilarity of the soil samples. The resultant NMDS shows that the soils with more diverse communities cluster away from those with simpler communities (Fig. 1A). The ordination first axis was strongly correlated with the Rock-Eval^®^ R-index (rho = −0.95, *P* < 0.0001) which quantifies the relative contribution of thermally stable compounds^7^. This is consistent with more diverse communities building more stable SOM^7^. Moreover, the bacterial community composition mirrored the SOM fingerprint (Fig. 1A, Procrustes statistics: 0.2070, *P* = 0.0057) indicating that the bacterial community composition drove the formation of SOM composition and quality.

**Figure 1.**
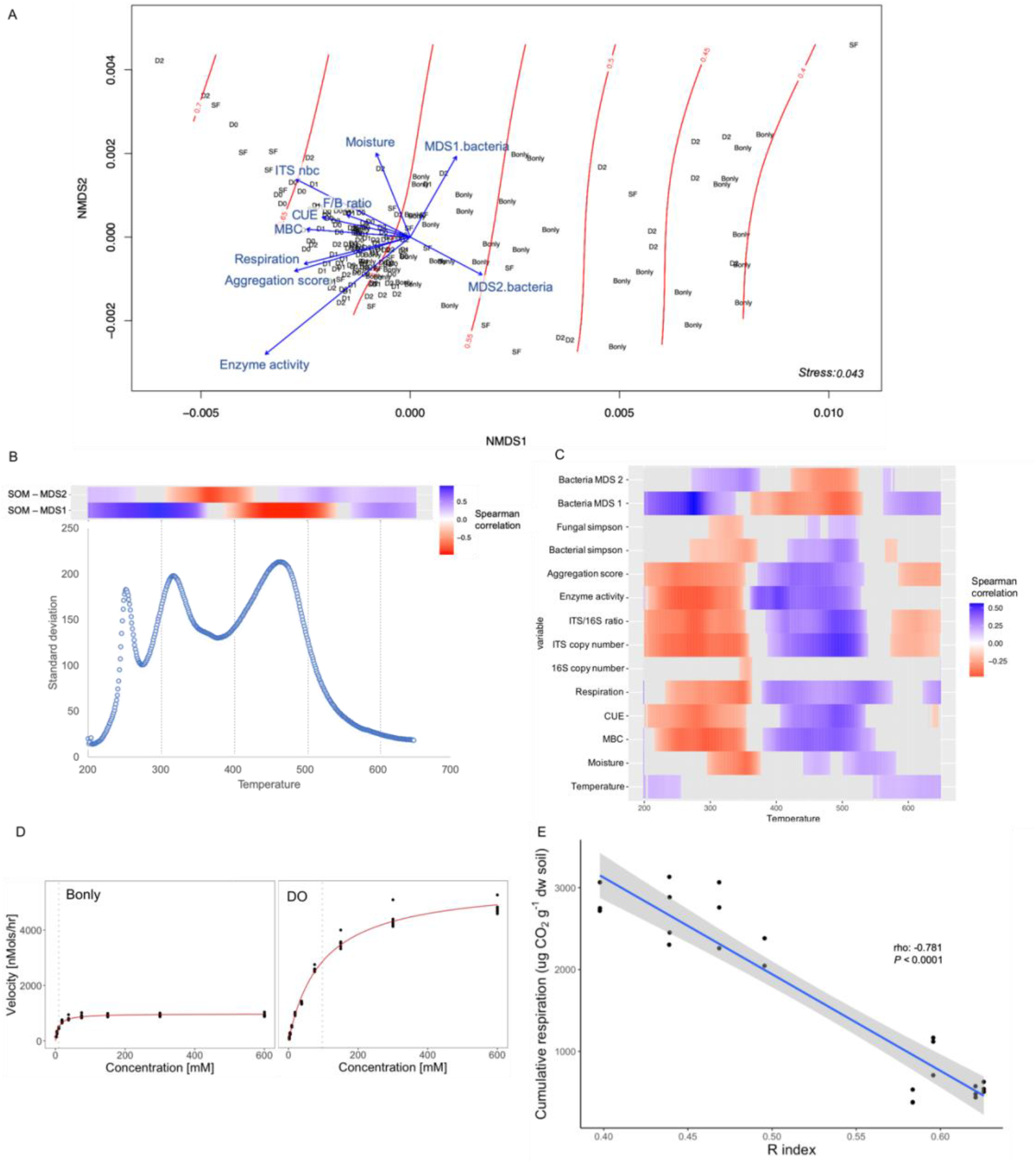
Drivers of SOM quality and stability. Ordination of soil organic matter quality. Non-metric multidimensional scaling of Bray Curtis distance from the pyrolyzed fraction of SOM based on Rock-Eval^®^ analysis. Red contour lines represent the SOM thermal-stability R index. Significant explanatory variables are represented as blue vectors (*P <* 0.05); enzyme activity corresponds to the maximum activity recorded (Vmax g^−1^ dw soil); MDS1- and MDS2.bacteria represent the first and second axis of the bacterial community structure, respectively; ITS and 16S copy number correspond to the quantification by qPCR of ITS and 16S rRNA gene (copy number g^−1^ dw soil); F/B ratio correspond to the fungal to bacterial ratio abundance; CUE represents the carbon use efficiency; MBC corresponds to microbial biomass carbon (µg C g^−1^ dw soil); Respiration represents the cumulative respiration measured during microcosms incubation (C-CO2 g^−1^ dw soil) and aggregation score represents the water stable aggregate formation at the end of incubation (A). Spearman correlation between the SOM ordinations axes points and the FID signal captured at different temperatures and standard deviation of signal across all microcosms by temperature (B). Spearman correlation between biotic variables and abiotic experimental treatment conditions and the FID signal captured at each temperature (C). Betaglucosidase enzymatic kinetics at representative samples for “Bonly” and DO treatments, vertical line represent the Km (D). The relationship between thermal-stability R-index and cumulative respiration (E).

Interestingly, the fungal community composition seemed to be less important in driving the SOM signature (Procrustes results; cor = 0.143, *P* = 0.0782). However, fungal abundance was positively related to the thermal stability of SOM (Fig. 1A and cor = 0.44, t = 6.4248, df = 168, *P* < 0.0001), supporting the role of fungi in overall community decomposition efficiency. This result is in agreement with research suggesting that fungi are major drivers of C cycling in soils^8^. Thus, while fungi were crucial for substrate decomposition, the SOM formed in these soils mirrored its bacterial community composition, suggesting that fungi and bacteria might play complementary roles during the decomposition and stabilization of new SOM.

We next looked in more detail the range of temperatures that captured the variation observed within the SOM ordination axes (Fig. 1B) and the biological community characteristics that drove the differences in the fingerprint of SOM (Fig. 1C). The signal captured at across the temperature range reflected differently to the microbial community biological characteristics. Bacterial community richness showed a positive saturating relationship with carbon use efficiency in these soils^4^ and our results here demonstrate that communities which grew more efficiently at the end of the incubation were also associated with more thermally-stable SOM (Fig. 1A-C). This is consistent with the theory stipulating that high growth efficiency is ultimately associated with greater soil carbon retention^9,10^. Communities depleted of fungi were characterized by low growth efficiency and low biomass in these soils, and also had some of the most thermally-labile SOM. Although low biomass was a predictor of the SOM signature (Fig. 1A-C), this was not due to residual simple added sugar. Indeed, the patterns of lower SOM thermostability in lower diversity communities held even when the lowest temperature, predominantly sugar-rich peak^11^, was removed (Supplementary material). Therefore, community composition acts independently of efficiency and biomass to drive SOM quality.

Using variance partitioning we investigated the drivers of SOM R-index (Supplementary material). Notably, microbial activity variables explained the most variance. This suggests that microbial activity and the by-products of their metabolism, such as extracellular enzymes, drove the formation of more stable SOM. This agrees with the idea that microbial processing of carbon contributes to the SOM pool^10^.

Fungi and bacteria are considered to play different roles in soil C cycling^12,13^. Accordingly, we observed distinct extracellular enzymatic dynamics in microcosms dominated by bacteria (“B_only_”) compared to microcosms with bacteria + fungi growing concomitantly. B_only_ microcosms showed a reduced maximum enzymatic activity (Fig. 1D) (V_max_) (*P* < 0.0001, F = 16.43, df = 136) and Michaelis constant (K_m_) (*P* < 0.01, F = 5.195, df = 136) compared to other dilution treatments which should result in a smaller uptake of C and reduced microbial turnover of SOM^9,12^. This could indicate that additional transformations of SOM occurred in the communities with greater complexity. Thus, the bacterial communities may have benefitted from by-products of fungal growth and metabolism^13,14^ - leading to increasingly thermally stable SOM. While previous findings suggest that decomposition of fungal residues is an important regulator of C accumulation is soils^8^, our results demonstrate that fungal ↔ bacterial interactions play an important role in this process.

Finally, to verify if more thermally stable SOM results in less available substrate C to microorganisms, in a subsequent experiment we inoculated a subset of microcosms with a diverse soil inoculum and measured cumulative respiration. We observed a negative correlation between thermal stability and cumulative respiration (Fig. 1E). This confirms that more thermally-stable carbon is less biodegradable and more likely to persist in the soil.

Model soils can be used to increase our understanding of major microbial ecology questions as it provides a single platform able to isolate specific components from confounding factors compared to natural soils^4,15^. Here, by using a model soil we show that microbial community composition and community characteristics drove the signature of the SOM and its thermal stability. Altogether, our results highlight the relevance of fungal ↔ bacterial interactions for the decomposition efficiency and the formation of new stable SOM.

## Materials and Methods

### Data collection

Microbial communities of varying complexities resulting from diluting and filtering soil were inoculated into an artificial soil matrix and amended with cellobiose and ammonium nitrate on a weekly basis^4^. At the end of four months of incubation, soil was characterized for respiration, growth, microbial biomass carbon, water-stable aggregate formation, betaglucosidase extracellular enzyme activity, fungal:bacterial phylogenetic marker gene copy ratio, and bacterial and fungal communities sequenced as described in Domeignoz-Horta et al. 2020^4^. Air-dried soil was subject to Rock-Eval^®^ ramped thermal pyrolysis following the protocol of Sebag et al. 2016^7^. To examine the relationship between SOM thermal stability and its bio-availability to microorganisms, we performed a follow up experiment in which we inoculated a natural soil-derived microbial inoculum into a subset of soils from the previous experiment. These microcosms were maintained at 60% water holding capacity and we monitored respiration daily during one week as a proxy for bio-available decomposable carbon.

## Data Analysis

Statistical analyses were performed in R statistical software (version 3.6.3), using the vegan and agricolae packages. For Rock-Eval^®^ analysis, soil samples are pyrolyzed from 200 to 650°C, followed by combustion of residual carbon from 400 to 850°C. Hydrocarbon compounds (HC) released during this process are measured through time by a flame ionization detector (FID) and generate a HC thermogram for each sample. Bray-Curtis distance of thermograms was calculated using mean C released at each 1°C unit of temperature and used as input to a non-metric multidimensional scaling (NMDS) to describe microcosms C signature. The variables that significantly explained the C signature amongst the different microcosms were identified using the *envfit* function (permutation tests n = 10000, *P* < 0.05). Surface fitting of variables within the ordination were performed using the *ordisurf* function. Here we used the thermogram generated during pyrolysis as a proxy for SOM chemical diversity. Procrustes analysis was used to test if the distance matrix of bacterial and fungal communities show superposition with the distance matrix of SOM (permutations test n = 10000, *P* < 0.05). We used Spearman correlations to explore relationships between C released at specific temperatures and the overall SOM ordination and biotic variables. We calculated the SOM thermal stability R-index as previously^7^.

## Data availability

The data and code supporting the findings presented here are available from the corresponding author on request and from Open Science Framework Repository project: https://osf.io/evb6d/.

## Acknowledgments

The authors thank Dr. Grace Pold for fruitful discussions. This work was supported in part by an award to KMD by the Department of Energy grant DE-SC0016590.

## Supplementary Materials

Variation partitioning analysis was applied to disentangle the contribution of biotic drivers and abiotic factors to the thermal stability R-index using the varpart function in the vegan package. Briefly, significant explanatory variables were selected using a backwards stepwise model selection process and an RDA (*P* < 0.05). Variables were then grouped into one of the following categories: microbial community diversity/structure, microbial activity, microbial abundance, or abiotic. All factors jointly explained 65% of the R-index variance, with activity being the strongest predictor contributing alone with 15%of the variance.

To verify that lower SOM signature was not driven by residual substrate we repeated our SOM ordination analysis excluding the signal captured in the first range of temperatures (200-275°C) as this is predominantly driven by a sugar-rich peak. This analysis showed very similar results to our analysis with the full dataset as the following variables significantly explained SOM ordination: moisture (r^2^ = 0.085, *P* = .0010), microbial biomass carbon (r^2^ = 0.12, *P* = .0002), carbon use efficiency (r^2^ = 0.087, *P* = .0014), bacteria MDS axis 1 (r^2^ = 0.095, *P* = .0005), bacteria MDS axis 2 (r^2^ = 0.0699, *P* = .0055), fungal:bacterial ratio (r^2^ = 0.048, *P* = .0457), aggregation score (r^2^ = 0.151, *P* < .0001), cumulative respiration (r^2^ = 0.151, *P* = .0001), ITS copy number (r^2^ = 0.187, *P* < .0001) and extracellular enzyme activity (Vmax) (r^2^ = 0.419, *P* < .0001).

## Notes

### Competing Interest Statement

The authors have declared no competing interest.

